# Genome-Wide Interrogation of SARS-CoV-2 RNA-Protein Interactions Uncovers Hidden Regulatory Sites

**DOI:** 10.1101/2025.05.26.656146

**Authors:** Joy S. Xiang, Karen X. Zhao, Laliv Tadri, Kurt Tamaru, Brian A. Yee, Katherine Rothamel, Jonathan C. Schmok, Jasmine R. Mueller, Samuel S. Park, Assael A. Madrigal, Rachael N. McVicar, Elizabeth M. Kwong, Ben A. Croker, Alex E. Clark, Aaron F. Carlin, Charly Acevedo, Declan McCole, Seán E. O’Leary, Rong Hai, Sandra L. Leibel, Gene W. Yeo

## Abstract

The global impact of the COVID-19 pandemic underscores the critical need for a comprehensive understanding of SARS-CoV-2 replication mechanisms. While the central roles of the RNA dependent RNA polymerase (NSP12), primase protein (NSP8), and nucleocapsid protein (N) in the virus life cycle are extensively studied, the precise nature of their interactions with the full-length viral RNA genome remain incompletely characterized. In this study, we sought to address this knowledge gap by employing enhanced crosslinking and immunoprecipitation (eCLIP) to map the binding sites of NSP8, NSP12, and N proteins across the SARS-CoV-2 genome at early stages of viral RNA and protein synthesis and late stages of virion assembly. Our findings revealed interactions of NSP8 and NSP12 to the 5’ and 3’ untranslated regions (UTRs) of both positive and negative sense RNA, regions known to regulate viral replication, transcription, and translation. We identified a surprising and essential NSP12 binding site within the RNA sequence encoding the conserved Y1 domain of NSP3, which regulates RNA abundance upstream of the site. Additionally, we found that N protein interacts with the 5’ UTR and influences translation efficiency. Finally, we report a novel regulatory function of N protein in modulating ribosomal frameshifting proximal to the frameshift element, a crucial process for maintaining viral protein stoichiometry. Our results provide a detailed molecular map of SARS-CoV-2 protein-RNA interactions, revealing potential therapeutic targets for attenuating viral fitness and informing the development of next-generation antiviral strategies.

## Introduction

COVID-19 is caused by the novel severe acute respiratory syndrome coronavirus 2 (SARS-CoV-2), a positive-sense single-stranded (+ss)RNA virus. The viral genome encodes 29 proteins^1^, which include the four structural proteins, membrane or matrix (M), nucleocapsid (N), envelope (E), and spike (S) proteins. In addition, there are 16 non-structural proteins, NSP1-16, and 9 accessory proteins ORF3a-ORF10, though evidence suggests the translation of several more peptides^2^. Replication, transcription and translation during the viral life cycle are regulated by complex mechanisms involving protein-RNA interactions that remain incompletely understood.

SARS-CoV-2 encodes replicase proteins that transcribe and replicate its positive sense virus genome, which also serves as an mRNA that initiates translation into proteins immediately upon entry into the cytoplasm. At the core of the replication-transcription complex is the RNA-dependent RNA polymerase (RdRp), also known as non-structural protein 12 (NSP12). NSP7 and NSP8 form the primase and stabilize RdRp for its activity^3^. RdRp replicates and transcribes viral RNAs and mRNAs with the help of the helicase NSP13, proofreading exonuclease NSP14, zinc finger protein NSP10 and ssRNA binding protein NSP9^4,5^. Viral RNA structures within the untranslated regions (UTR) facilitate the assembly of the primase and RdRp to initiate transcription and replication. The 5′ UTR consists of conserved RNA stem loop structures 1–4 (SL1-4), while stem loop 5 (SL5) extends partway into the open reading frame of the first gene *ORF1ab*. Stem loops 1–3 have been linked to the regulation of coronavirus replication^6,7^ and transcription of subgenomic RNA^8^. The 3′ UTR contains a pseudoknot structure that binds the NSP7/NSP8 primase complex to initiate viral replication from the 3′ end^9^. Despite the importance of these structures, our understanding of the precise interplay between viral proteins and RNAs is limited.

Similar to transcription and replication, SARS-CoV-2 translation is intricately regulated. Stem Loop 5, which harbors the AUG start codon of the main open reading frame ORF1ab, is a highly conserved and stable structure in the 5′ UTR that is essential for the overall architecture of the 5′ UTR^10^. Stem Loop 4 contains an AUG that is the start of an upstream open reading frame (uORF) which ends before Stem Loop 5. Since uORFs are typically associated with suppressing translation initiation of the main open reading frame, SL4 may add another layer of translation regulation. Conserved among many viruses, SARS-CoV-2 encodes an essential frameshift element that causes ribosome slippage to shift the translation reading frame by one nucleotide in the 5′ direction. This is controlled by multiple conformations of RNA structures, an attenuator hairpin and a pseudoknot^11,12^.

While the role of NSP12 in replication and transcription is well established, recent evidence reveals its role in RNA capping^5,13^, expanding our understanding of its role in translation. Evidence for NSP12 interacting with translation elongation and regulatory factors eEF1A, UBAP2 and UBAP2L has also been found^14^. Both immediately after infection and at later stages, the nucleocapsid protein N binds viral genomic RNA to facilitate genome assembly and coordinates with the matrix protein M for virion packaging and maturation. In bovine coronavirus (BCoV), N protein binding to the viral poly(A) tail has been associated with translational inhibition^15^, although it remains unclear whether this effect is due to poly(A) binding itself or indirect interactions with translation initiation factors such as eIF4G/eIF4F. While suggestive, the relevance of this mechanism to SARS-CoV-2 translation remains uncertain. Together, these observations point to a more complex interplay of viral RNA with these essential viral proteins. Nevertheless, our knowledge of the genome-wide RNA-protein interaction in authentic virus infection remains lacking.

In this study, we comprehensively interrogate the direct interaction of SARS-CoV-2 proteins with virus RNA using enhanced crosslinking and immunoprecipitation (eCLIP^16,17^) in the context of authentic virus infection. eCLIP is a powerful tool which has been used to profile more than 150 human RBPs and provide insights into their regulatory roles^18^. We aim to identify interaction sites between NSP8, NSP12 and N with virus RNA that regulate mRNA and protein expression and elucidate their role in viral fitness.

## Results

### eCLIP elucidates SARS-CoV-2 protein-viral RNA interactions in virus infected cells

Due to their critical roles in replication, transcription and genome assembly, we chose to profile the interaction of viral RNA with SARS-CoV-2 primase protein NSP8, RdRp NSP12, and N. We performed eCLIP^16^ on SARS-CoV-2 infected African Green Monkey kidney (Vero E6) cells, which are an efficiently infected cell line **(Fig. 1a**). To assess interactions at early and late stages of infection, cells were infected at a high MOI (1) for 6 hours to ensure sufficient viral RNA early in infection, and at a low MOI (0.01) for 48 hours to allow extended infection without excessive cytopathic effects. Infected cells were subject to UV irradiation, which covalently crosslinked interacting proteins to RNAs. This was followed by immunoprecipitation using protein-specific antibodies to isolate the RNA bound to each protein. The RNA-bound proteins were resolved via SDS-PAGE and transferred to nitrocellulose membranes, which bind and retain the protein-bound RNA. The region spanning the expected protein size and 75 kDa larger were excised and purified in subsequent steps. The same size region of a non-immunoprecipitated input whole cell lysate was included as size-matched input (SMInput, or IN) to identify background crosslinked RNAs in the cell milieu (**Supplementary Fig. 1**). RNA was converted to DNA libraries and sequenced to an average depth of ∼25 million reads. The sequenced reads were mapped to the SARS-CoV-2 viral genome to determine SARS-CoV-2 protein RNA interactions.

**Fig. 1.**
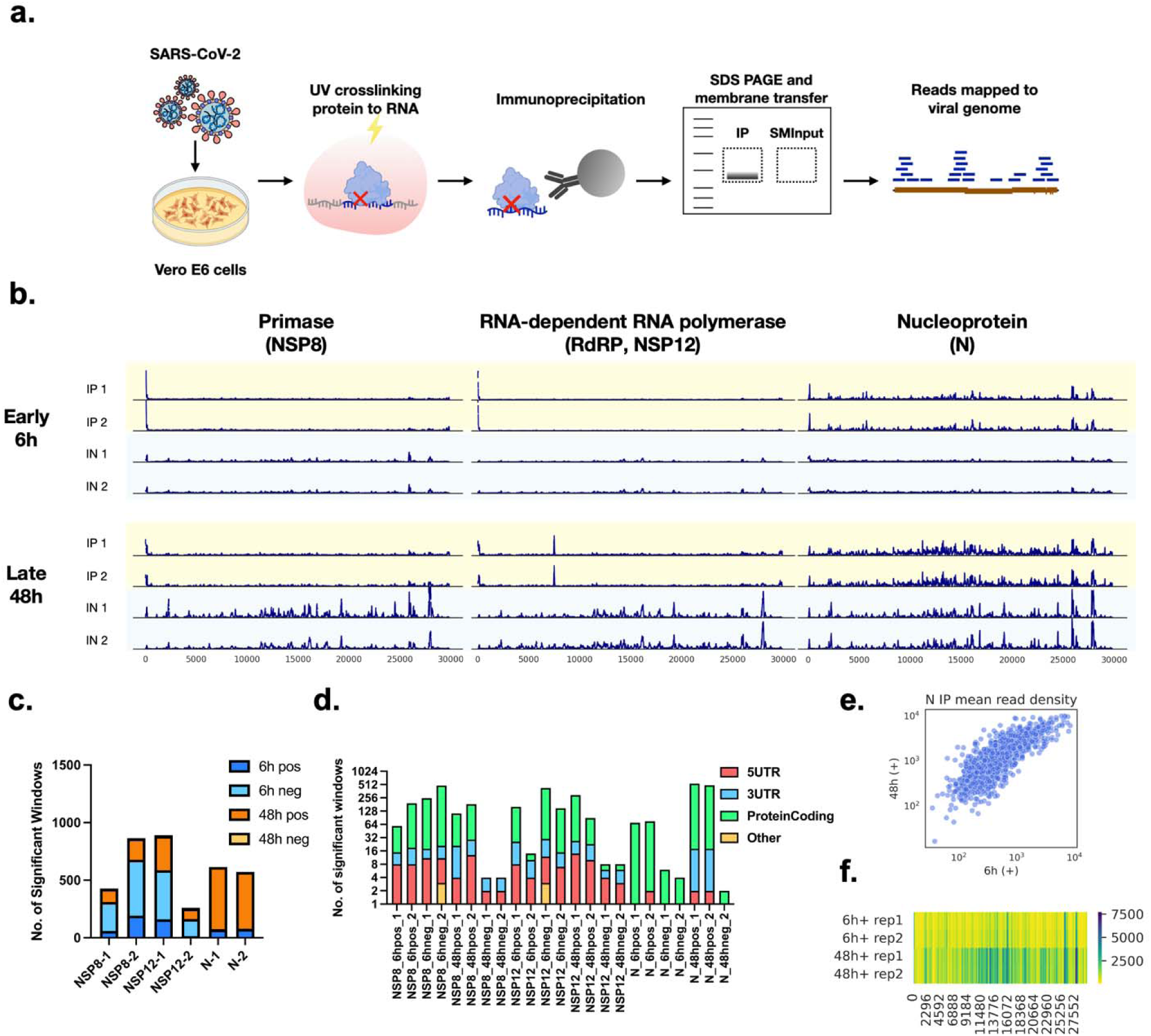
eCLIP maps SARS-CoV-2 viral RNA interactions with NSP8, NSP12 and N. **a)** Enhanced crosslinking and immunoprecipitation resolves RNA bound to SARS-CoV-2 proteins in virus infected cells. After SDS-PAGE and membrane transfer, the region corresponding to the Immunoprecipitated (IP) protein and 75kDa above is excised for RNA extraction. Size-matched input (IN) is the same region in the cell lysate to determine background crosslinked RNA. **b)** Read density plots show RNA enriched by N, NSP8 and NSP12, distributed across the SARS-CoV-2 genome on the positive strand. Two biological replicates were performed for each sample and condition. **c)** Bar plot indicating the number of significantly enriched 20 nucleotide (nt) windows of IP vs IN samples for each replicate sample and condition. Significance, log2foldchange>1.5, qvalue<0.01. **d)** Bar plot showing the breakdown of the number of significant windows in each region. 5UTR, 5′ untranslated region; 3UTR, 3′ untranslated region. **e)** Scatter plot showing the mean read density of each 20 nt window across the positive sense virus genome in N IP samples at 6h and 48h. **f)** Heatmap showing the read density of N IP samples (colorbar) as 200 nt sliding window mean across the SARS-CoV-2 genome.

Our eCLIP results provide a genome-wide map of RNA interactions with viral proteins during authentic SARS-CoV-2 infection (**Fig. 1b, Supplementary Fig. 2**). Reads are mapped to the virus genome and analyzed in windows of 20 nucleotides (**Supplementary Data**). Differences in read densities in each window of immunoprecipitated (IP) and input (IN) samples are compared to determine the fold enrichment and significance (two-tailed unpaired t-test, log2foldchange >1.5, q-value < 0.01). At 6 hours post-infection, both NSP8 and NSP12 show a greater number of significantly enriched windows on the negative-sense RNA compared to the positive-sense RNA (**Fig. 1c**). By contrast, at 48 hours post-infection, few such enrichments are observed on the negative strand. Notably, both NSP8 and NSP12 bind to regions in the 5′ and 3′ untranslated regions (UTRs) of both strands, which are known to regulate transcription and replication. These patterns indicate that NSP8 and NSP12 interact with both the positive- and negative-sense RNA during the early stages of infection, when transcription and replication are most active (**Fig. 1d**).

The N protein enriches few or no significant windows on the negative strand at both early and late time points (**Fig. 1c**). At 6 hours post-infection, significantly enriched windows in N eCLIP are almost exclusively within protein-coding regions, indicating a minimal role in regulating transcription and replication. However, by 48 hours post-infection, significantly enriched windows are distributed across most regions, agreeing with our conventional understanding that N protein interacts with the whole viral genome during late infection (**Fig. 1d**). N protein exhibits the greatest number of significantly enriched windows on the positive strand at 48 hours post-infection, when most replication is complete, and N protein is condensing the viral genome to facilitate packaging of the positive-strand genome into virions. Notably, in the immunoprecipitated samples, regions with high read density at 6 hours remain enriched at 48 hours, while additional regions of the genome increasingly come into contact with N. (**Fig. 1e-f**). Our observations align with our understanding of the role of N in genome RNA association for virus assembly^19^. The increase in the number of interactions could be due to a combination of increasing concentrations of N available to bind to weaker sites and time needed for genome assembly and condensation to take place. All in all, our eCLIP data provide a global view of viral RNA interactions with the three essential proteins involved in critical processes of the viral life cycle.

### 5′ UTR and 3′ UTR regions of both positive and negative strands interact with NSP8 and NSP12

At the early stage of infection (6h post-infection), significantly enriched read densities in NSP8 and NSP12 samples are primarily clustered at the 5′ untranslated region (5′ UTR) of the positive sense strand (**Fig. 2a**). The 5′ UTR is a highly structured region known to regulate replication and transcription^20^. Therefore, 5′ UTR interaction with replicase proteins NSP8 and NSP12 is expected. Specifically, NSP8 and NSP12 eCLIP reads cover the first three stem loop structures (SL1-SL3). The mean fold enrichment of NSP8 and NSP12 are 25 (p<1e-28) and 28 (p<1e-20) in the first 75 nucleotides (nt), respectively (**Fig. 2b**). SL1 and SL2 are known to regulate replication^6,7^, and SL3 contains the leader transcription regulating sequence (TRS-L) responsible for the discontinuous transcription of subgenomic mRNAs facilitated by NSP12 template switching^21^. The enrichment of reads at the 5′ end likely indicates a rate-limiting step in initiating replication-transcription due to the slow assembly of the NSP12–NSP7–NSP8–RNA core replicase complex^22,23^. The high read density could also be due to the accumulation of replicase at the 3′ end of the negative strand after completing the synthesis of the negative strand before re-initiating synthesis of the positive strand.

**Fig 2.**
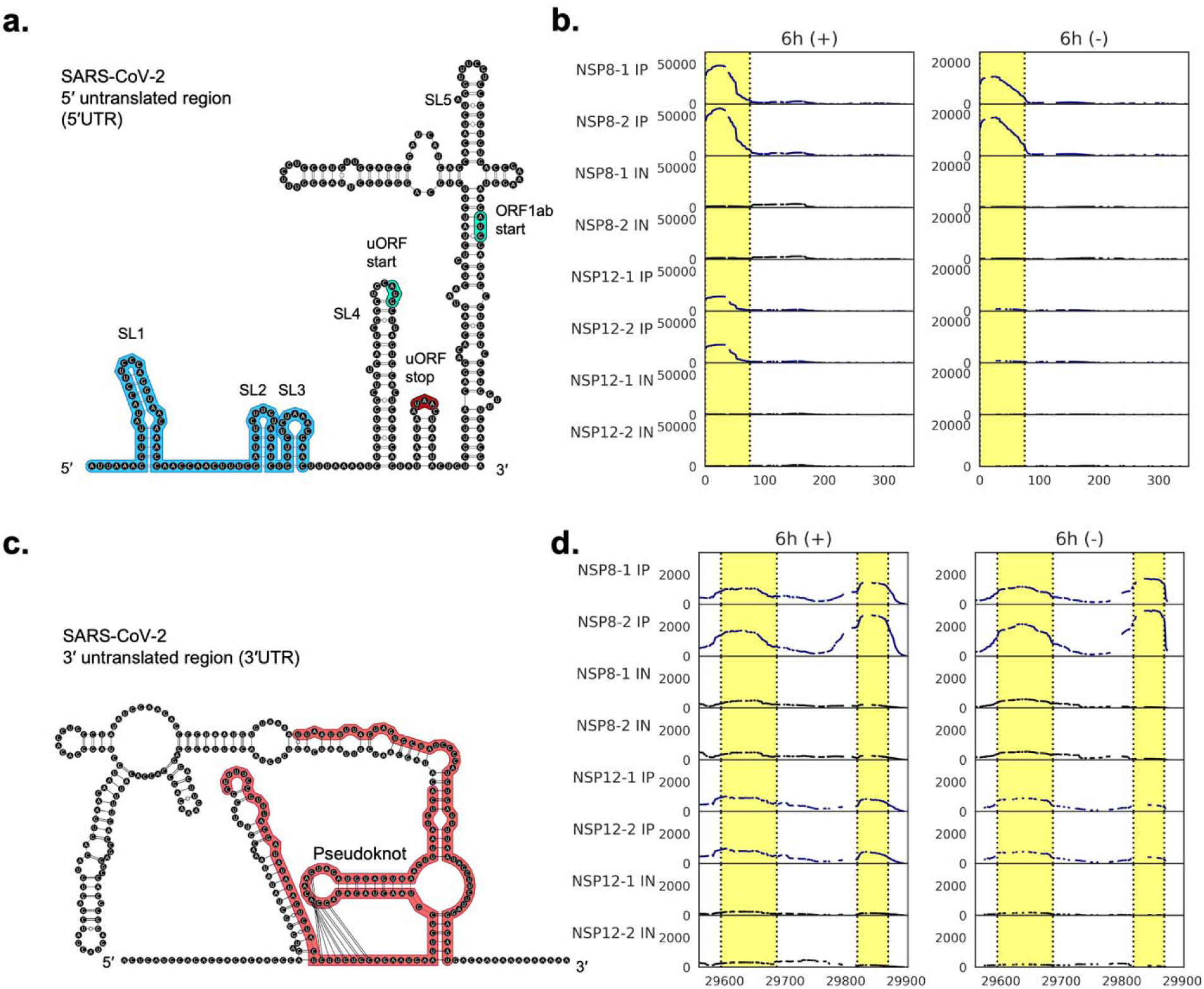
eCLIP highlights interactions between replicase proteins NSP8 and NSP12 at the 5′ and 3′ untranslated regions. **a)** Secondary structure of the region from the 5′ untranslated region to the end of the conserved stem loop 5 (SL5). Genome coordinates use positive strand numbering. Highlighted in blue are regions bound by NSP8 and NSP12, corresponding to yellow highlighted regions in **b**. Conserved stem loop structures (SLs), start codons (green) and stop codon (maroon) are annotated for the upper open reading frame (uORF) and the beginning of *ORF1ab*. Structure is predicted using RNAfold. **b)** Read density plots showing RNA enriched by NSP8 and NSP12 eCLIP on the positive (left) and negative (right) strand near the 5′ end of the positive sense genome. **c)** Secondary structure of the region from the N protein stop codon to the 3′ end of the positive strand. Highlighted in red are regions bound by NSP8 and NSP12, corresponding with yellow highlighted regions in **d.** Structure predicted using RNAfold and pseudoknot annotation from ref^41^. **d)** Read density plots showing RNA enriched by NSP8 and NSP12 on the positive (left) and negative (right) strand near the 3′ end of the positive sense genome.

On the negative strand, NSP8 enriched significant windows are clustered at the 3′ end, which corresponds to the 5′ end of the positive-sense genome. The mean fold enrichment of NSP8 is 39 (p<1e-41) in the 0-75 nucleotide region, compared to 9 for NSP12 (p<1e-38). The mean read density of NSP12 IP samples is 765, while that of NSP8 IP samples is 11,454, suggesting NSP8 plays a significantly greater role in interacting with the 3′ end of the negative strand. These enrichments may not necessarily represent separate binding events to free positive- and negative-sense RNAs. Instead, the observed binding to both strands at similar regions may reflect interactions with duplex RNA, as the replicase complex, which comprises NSP8 and NSP12, transits the 5′ end of the positive strand during transcription elongation, or switches from the 3′ end of the negative strand to the positive strand during replication.

At the 3′ end of the positive strand are conserved stem loop structures known to regulate replication. Among them is a pseudoknot structure that recruits primase proteins NSP7 and NSP8, which as a complex synthesizes the first nucleotides of a short primer that is elongated by the RNA polymerase NSP12 for replication and transcription^11,24^. Our eCLIP data provides evidence of direct binding interaction at the 3′ untranslated region (3′ UTR) in support of this model understanding of replication-transcription initiation (**Fig. 2c**). The mean fold enrichment of NSP8 is 2.9 (p<1e-37) at region 29,594–29,686 nt and 9.6 (p<1e-51) at region 29,819–29,870 nt. NSP12 similarly enriches at these two regions, with fold enrichment of 3.4 (p<1e-51) at region 29,594-29,686 nt and 5.5 (p<1e-33) at region 29,819-29,870 nt. Similar to the 5′ end, we observed significant enrichment on the negative strand in the complementary regions. The mean fold enrichment of NSP8 is 4.6 (p<1e-52) at region 29,594-29,686 nt and 10 (p<1e-18) at region 29,819-29,870 nt. NSP12 IP reads are similarly enriched at these two regions, with fold enrichment of 2.9 (p<1e-42) at region 29,594-29,686 nt and 41 (p<1e-60) at region 29,819-29,870 nt (**Fig. 2d**). While this mirroring interaction on the negative strand is expected, we previously lacked direct evidence to confirm it. The NSP7/NSP8 and NSP12 RdRp complex together has a large RNA-binding interface, and this is reflected by the many nucleotides that contact both NSP12 and NSP8 at the 3′ end structure^25^.

### Novel interaction between RdRp NSP12 and the RNA encoding the conserved Y1 domain of NSP3

As the RNA-dependent RNA polymerase (RdRp) of SARS-CoV-2, NSP12’s primary role is in replicating and transcribing viral RNA and contributing to the cap formation of the mRNA. Rate limiting steps for these reactions are at the 5′ and 3′ UTRs, corresponding to enriched eCLIP reads at these regions at early stages of infection. Therefore, we were surprised to find that NSP12 binds to an RNA sequence within the coding region of NSP3 (**Fig. 3a**). We observed that the peak in read density at positions 7436–7526 is uniquely enriched in NSP12, with a mean fold enrichment of 5.1 (p < 1e-23) at 48 hours post-infection, predominantly on the positive strand, and a fold enrichment of 3.6 (p < 1e-16) on the negative strand at 6 hours post-infection (**Fig. 3b**). While there is no significant enrichment on the positive strand at 6h post infection, there is a distinct peak in the IP samples. This enriched region is an RNA sequence encoding an annotated Y1 domain, which is highly conserved among nidoviruses, while the region after (CoV Y) is conserved in all coronaviruses^26^. To verify, we performed a phylogenetic analysis of SARS-like betacoronaviruses and multiple sequence alignment of their genomes (**Fig. 3c**). From this alignment, we determined the Shannon entropy of each nucleotide sequence across the genomes, using SARS-CoV-2 as a reference (**Fig. 3d**). The NSP12 footprint is located at a region of low entropy, indicating that it is relatively conserved. Even though we know very little about the function of Y1 + CoV Y, the conservation of the Y1 domain and the strong enrichment of this region bound by NSP12 led us to hypothesize that the interaction plays a crucial role in viral fitness.

**Fig 3.**
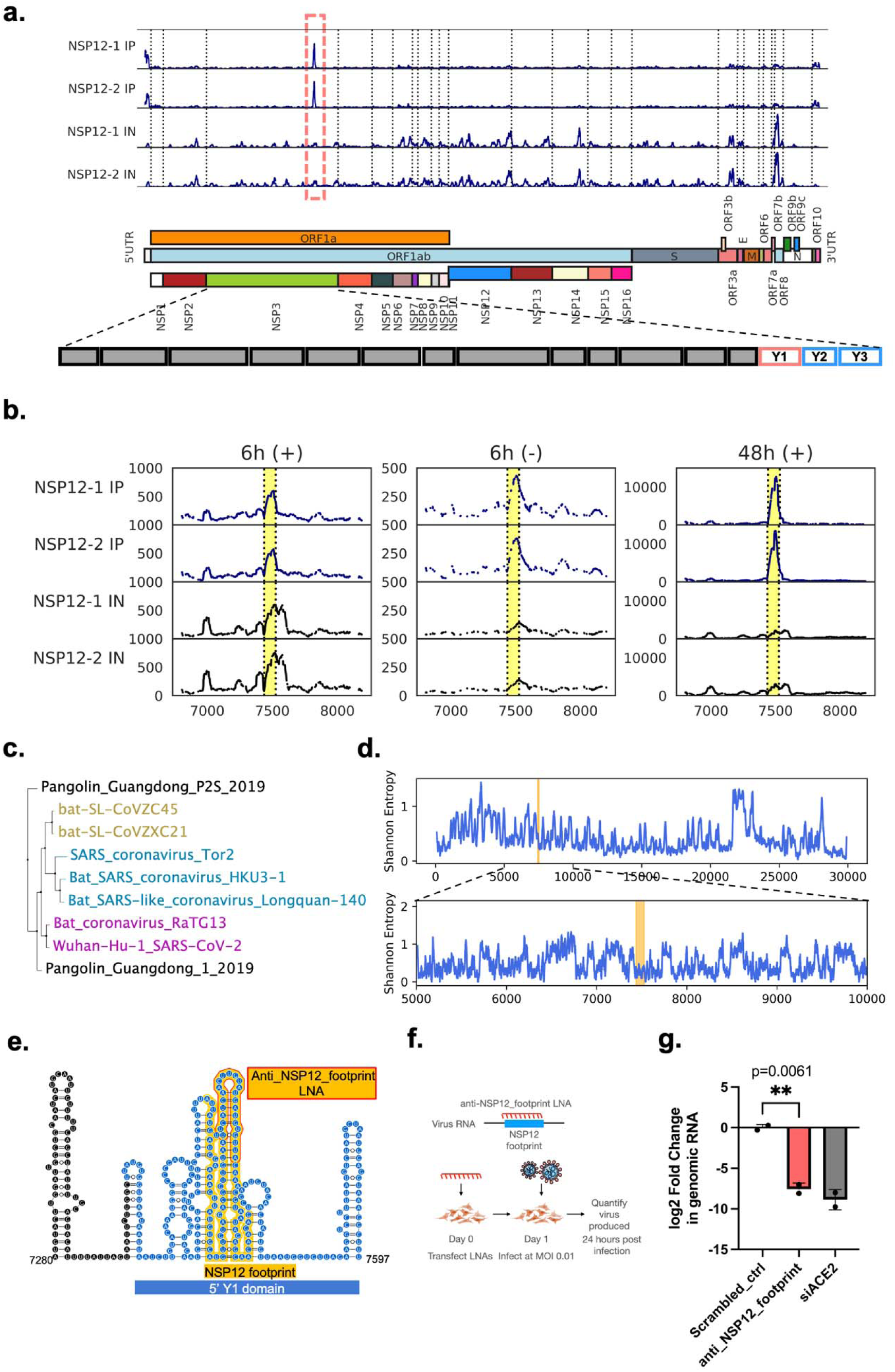
NSP12 binds to an RNA sequence within the conserved NSP3 Y1 domain coding sequence. **a)** Read density plot highlighting the peak that shows maximum read density across the genome in the NSP12 IP samples. Black dotted lines indicate translation start of annotated open reading frames and the start of each non-structural protein, corresponding to the schematic of SARS-CoV-2 genome. Orange dashed line box zooms into the bound region. Genome annotation indicates the bound region is in NSP3. NSP3 domains are illustrated in a schematic with different grey boxes, the Y1 domain coding sequence is outlined in orange, to which NSP12 binds. Y1+Y2+Y3 form the Y1+CoV Y domain conserved among coronaviruses. **b)** Zoomed in read density plots indicating enrichment of NSP12 bound RNA at the region 7436 – 7526, marked by dotted lines. **c)** Conservation analysis determined from an alignment of 9 coronavirus species using MAFFT v7.453 (2019/Nov/8). The phylogenetic tree is constructed based on the alignment using a neighbor joining algorithm in JalView version 2.11.4.1. **d)** The aligned sequences in (c) were used to calculate the Shannon entropy of nucleotide identity at each genomic position. Top: Shannon entropy across the entire SARS-CoV-2 genome using a sliding window mean of 100 nucleotides. Bottom: Shannon entropy focused on the region near NSP3 Y1 domain; the orange-highlighted region corresponds to the 7436–7526 nt region bound by NSP12. **e)** RNA secondary structure of the region around the NSP12 footprint obtained from ref^11^. **f)** Schematic showing the LNA treatment to test the importance of the NSP12 footprint to virus replication. **g)** RT-qPCR results indicating log2 fold change of SARS-CoV-2 genomic RNA levels in virions released from cells treated with anti_NSP12_footprint LNA, compared to a scrambled LNA control and siRNAs targeting ACE2. * two tailed t-test.

To determine if the Y1 coding sequence contributes to virus proliferation, we treated cells with an antisense locked nucleic acid (LNA) – anti-NSP12_footprint – designed to target the predicted RNA stem loop within the NSP12 footprint region (**Fig. 3e**). Treating SARS-CoV-2-infected cells with LNAs targeting structured regions is an established assay for assessing the importance of viral RNA structures to virus fitness^11^. This is achieved by disrupting RNA structures and/or interactions with RNA-binding proteins through base pairing with complementary LNAs. The anti-NSP12_footprint LNA was designed with a melting temperature similar to or lower than those of previously reported LNAs^11^ to minimize the likelihood of blocking polymerase elongation or translation (**Supplementary Table 1**). We treated cells with the anti-NSP12_footprint LNA and a scrambled control LNA^11^ and quantified virus production by measuring genomic RNA from virions released by the treated cells. We observed a significant reduction in the amount of virus produced (**Fig. 3f-g**) by cells treated with the anti-NSP12_footprint LNA compared to the scrambled LNA control. Correspondingly, the number of plaques produced and the amount of virus protein and RNA within the cells are reduced, supporting the data that the LNA treatment impeded virus infection and replication (**Supplementary Fig. 3**). No cytotoxicity was observed compared to the scrambled LNA control (**Supplementary Fig. 4**). For comparison, we designed an adjacent anti-NSP12_footprint_neg LNA which showed minimal/no effect on virus levels (**Supplementary Fig. 3c**), indicating that our conservative melting temperature design likely biases towards false negatives. Our results thus suggest that the Y1 NSP12 footprint region partners with NSP12 to play an important role in the virus life cycle.

### NSP12 differentially regulates RNA levels upstream of the binding site

We hypothesize that NSP12 binding to the Y1 coding region may influence RNA abundance or the expression of proteins upstream or downstream of the binding site. The enriched read density of NSP12 at this footprint could result from a slowdown in polymerase elongation, a non-replicative binding mode, or both. Consequently, any RNA or protein expression regulation could occur through multiple mechanisms, including premature termination or alternative initiation of polymerase activity, or modulation of RNA stability or translation efficiency. To begin exploring this, we developed a reporter system where *Cypridinia* luciferase (CLuc) and *Gaussia* luciferase (GLuc) are linked by the 5′ start of the Y1 coding sequence up to the end of the NSP12 footprint region (**Fig. 4a**). This is to mimic the long open reading frame *ORF1ab* that contains the footprint region. As a control, we scrambled the DNA sequence of the 5′ Y1 and NSP12 footprint regions without changing the amino acid sequence i.e. synonymous mutations, to avoid any expression differences that could arise from changing the peptide linker. This allows us to test the RNA sequence-dependent effect of the NSP12 footprint on regulating protein expression and its interaction with NSP12. We found that NSP12 significantly increased the luciferase activity of CLuc when the NSP12 footprint is present. Scrambling the 5′ Y1 sequence but having an intact NSP12 footprint led to similar levels of induction (**Fig. 4b**). Scrambling the NSP12 footprint abolished the induction effect on CLuc activity by NSP12. In contrast to the CLuc data, GLuc activities are the same between NSP12 and mCherry expression across all four reporters (**Fig. 4c**). Accordingly, the ratio of CLuc to GLuc increases upon NSP12 overexpression (**Fig. 4d**). This differential regulatory effect on luciferase activity is also observed at the RNA level of CLuc relative to GLuc. (**Fig. 4e**). Intriguingly, the NSP12 footprint region lies about 0.5-1.5 kilobases downstream of a steep, around ten-fold drop off in SARS-CoV-2 RNA reads in both the positive and negative strand^27,28^ (**Fig. 4f**). While the mechanism underlying this abrupt change in RNA levels remains unknown, it raises the possibility that NSP12 binding to this region may be one contributing factor.

**Fig 4.**
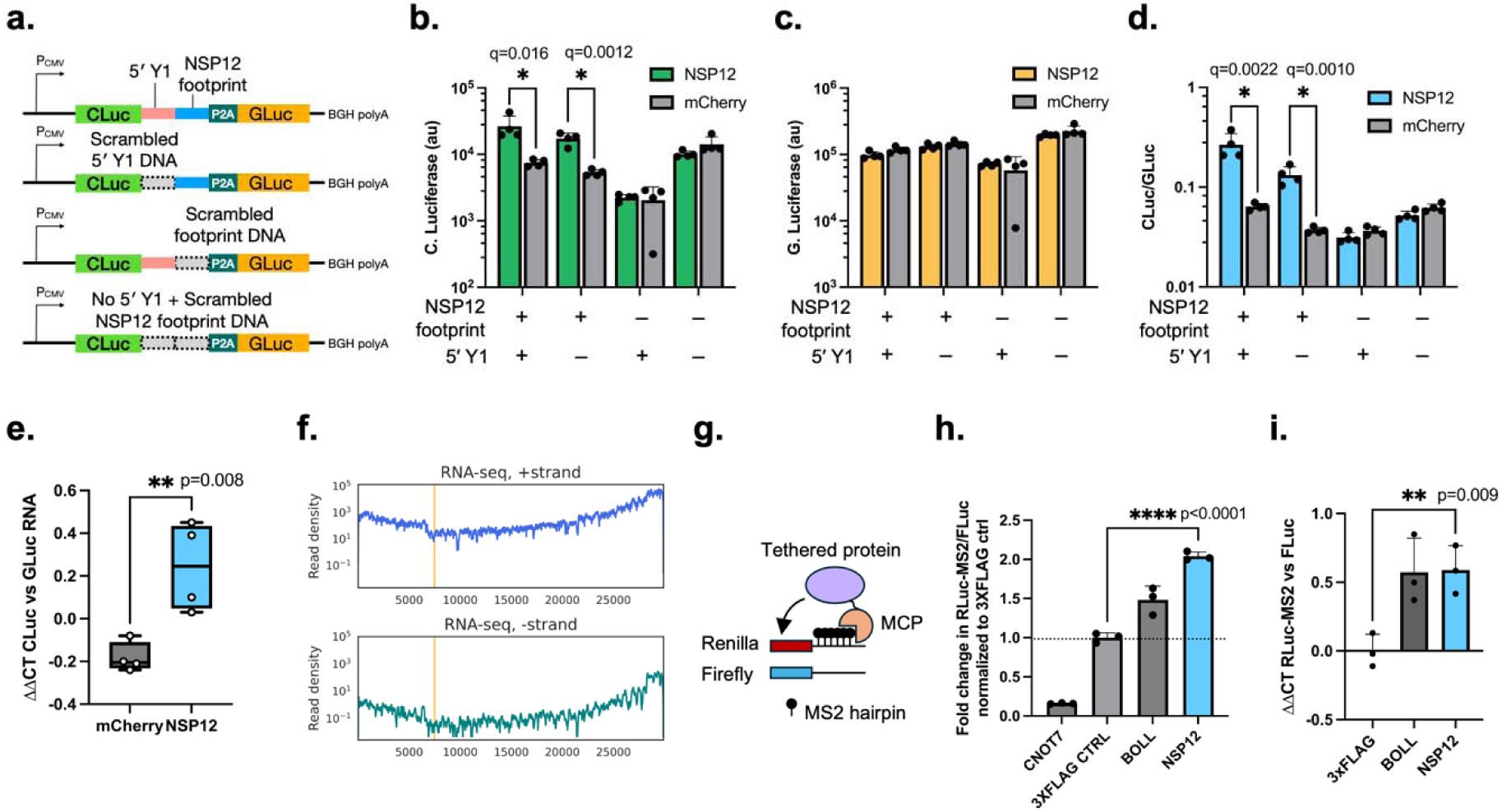
NSP12 differentially regulates RNA levels upstream of its binding site. **a)** Schematic showing a dual luciferase reporter assay where 5’ Y1 + NSP12 footprint sequences are inserted in between Cypridinia and Gaussia luciferases, CLuc and Gluc respectively. As negative controls, the DNA sequences are scrambled while preserving the amino acid sequence. **b-d**) Luciferase activity of CLuc (**b**), GLuc (**c**), and ratio of CLuc vs. GLuc (**d**) when reporters in (**a**) are co-transfected with NSP12 and mCherry negative control. q value is computed from multiple two-tailed unpaired t-test using Benjamini-Hochberg correction. **e)** Ratio of *CLuc* to *GLuc* RNA levels determined by RT-qPCR. ΔΔCT, log2 fold difference of *CLuc* RNA compared to *GLuc* RNA. **f)** PolyA mRNA-seq data showing the abundance of virus RNA sequencing read density across the genome. The orange line indicates the NSP12 footprint region 7436 – 7526. **g)** Schematic representation of a dual luciferase reporter assay utilizing two plasmids encoding Renilla luciferase (RLuc) and Firefly luciferase (FLuc). Six MS2 RNA hairpins are integrated into the 3′ untranslated region (UTR) of RLuc, enabling binding by the MS2 phage coat protein (MCP). A regulatory protein that modulates RNA stability or translation efficiency is fused to MCP, facilitating its recruitment to RLuc via the hairpins. The resulting changes in RLuc expression are quantified relative to the FLuc transfection control. **h)** Fold change in the RLuc/FLuc luciferase activity ratio for MCP-tagged proteins. 3×FLAG serves as a non-regulatory control, CNOT7 as a downregulatory control, and BOLL as an upregulatory control. **i)** Fold change in the *RLuc*/*FLuc* RNA ratio in an experiment analogous to (**h**). All p-values are determined by two-tailed unpaired t-tests.

To verify that NSP12 has a stabilizing effect on upstream RNA, we employed a different dual luciferase assay previously reported to screen around 700 human RNA binding proteins for their regulatory function^29^. One reporter plasmid encodes a *Renilla* luciferase (RLuc) with six MS2 phage RNA hairpins inserted in the 3′ UTR. A transfection control plasmid encodes *Firefly* luciferase (FLuc) without any MS2 hairpins. NSP12 is fused to the MS2 phage coat protein (MCP), which binds the RNA hairpins with high affinity, recruiting NSP12 to the 3′ UTR of RLuc mRNA (**Fig. 4g**). CNOT7 fused with MCP serves as a downregulatory RNA degradation control, BOLL fused with MCP serves as an upregulatory translation-enhancing control, and a 3×FLAG peptide similarly fused to MCP serves as a non-regulatory negative control. By determining the ratio of RLuc to Fluc luciferase activity, we found that NSP12 significantly increases the targeted RLuc expression (**Fig. 4h**). This increase is also observed at the RNA level, as reflected in the ratio of RLuc to FLuc (**Fig. 4i**). Together with the above-mentioned RNA-seq data from SARS-CoV-2-infected cells, we conclude that NSP12 increases the RNA level of the sequence that comes before its binding site, which may also increase upstream protein production. However, we are only at the beginning of understanding the functional consequences of this particular NSP12-RNA interaction. Alternative interpretations, which include indirect or coincidental effects, remain plausible. Further studies will be needed to determine whether NSP12 is directly involved in modulating RNA abundance, and if so, by which mechanism.

### N protein binds to the 5**′** UTR stem loop 5

While significantly enriched windows in N eCLIP are mostly distributed throughout the genome with few windows in the untranslated regions, we were surprised to find a region in the 5′ UTR at 6h post infection with the highest read density in the IP samples (**Fig. 5a-b**). From position 139 to 181, the mean fold enrichment of IP to IN is 2.5 (p<1e-18) (**Fig. 5b**). The same region is enriched at late infection, though to a lesser extent. This region is directly downstream of the stop codon of an upstream open reading frame (uORF) conserved in SARS-CoV-2 variants^11,20,21^. The bound region extends into the 5′ end of the stem loop 5 (SL5, at position 150 to 294 nucleotides), a large 4-way junction structure highly conserved across coronaviruses but with largely unknown function^30^. Since a high level of virus translation occurs at early stages of virus infection, we hypothesized that N’s association with the 5′ UTR at an early infection stage suggests a potential role in regulating translation initiation. Additionally, Shannon entropy analysis indicates that the N footprint region is conserved among related betacoronaviruses (**Fig. 5c**), which led us to hypothesize that it plays an important role in the virus life cycle.

**Fig 5.**
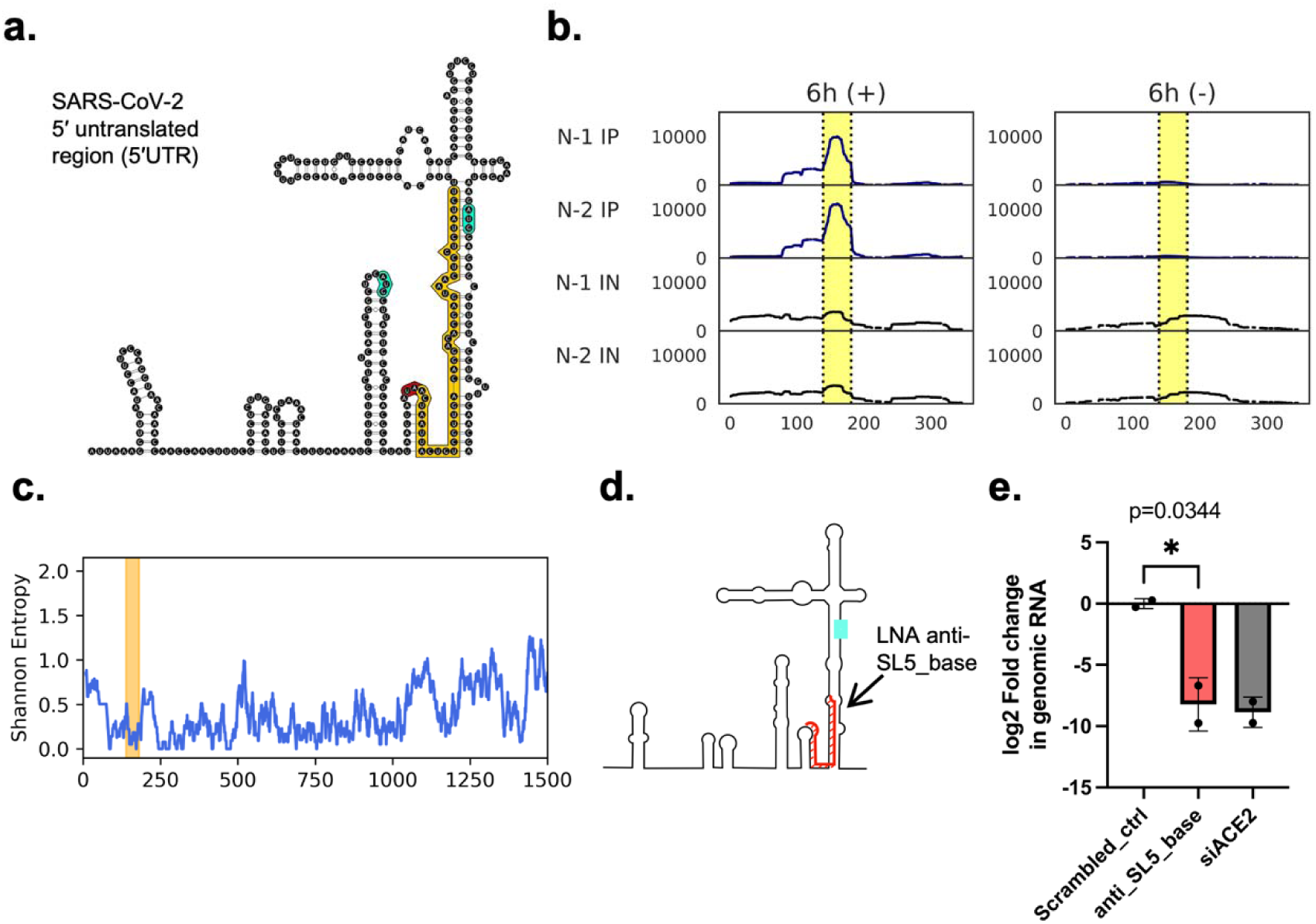
N interacts with the 5′ base of stem loop 5 in the 5′ untranslated region. **a)** Secondary structure from the 5′ untranslated region (UTR) to the end of the conserved stem loop 5 (SL5). Highlighted in yellow is a region that interacts with N corresponding to the highlighted regions in **b**. **b)** Read density plots showing RNA enriched by N eCLIP near the 5′ start of SARS-CoV-2 RNA. **c)** Shannon entropy conservation analysis focused on the 5′ region of SARS-CoV-2 RNA; the orange-highlighted region corresponds to the 139 – 181 nt region bound by N. **d)** Schematic showing the region targeted by a locked nucleic acid (LNA) complementary to the N interaction site. **e)** RT-qPCR results indicating log2 fold change of SARS-CoV-2 genomic RNA levels in virions released from cells treated with anti_SL5_base LNA, compared to a scrambled LNA control and siRNAs targeting ACE2. *p-values are from two-tailed unpaired t-tests.

To determine whether the region bound by N is essential to the virus, we designed an LNA – anti-SL5_base – that targets the 5′ proximal end (141 – 161 nt) bound by N (**Fig. 5d**). We hypothesized that disrupting interactions between this region and regulatory factors, such as N, would affect viral proliferation. The LNA was designed with a similarly conservative melting temperature as the anti-NSP12_footprint LNA (described in the previous section) to minimize the risk of interfering with the translation initiation scanning complex. We treated cells with the anti-SL5_base LNA and a scrambled control LNA, and observed a significant reduction in virus production in cells treated with anti-SL5_base LNA (**Fig. 5e, Supplementary Fig. 3**), with no noticeable cytotoxicity compared to the scrambled control (**Supplementary Fig. 4**). As a negative comparison, an adjacent LNA (anti-anti-AUG) that targets the region complementary to the *ORF1ab* AUG codon had no significant effect on virus RNA levels (**Supplementary Fig. 3d**). These results indicate that the N footprint at the 5′ base of SL5 serves as an important regulatory site for viral replication.

### N regulates translation via the 5**′** UTR binding site

Next, we developed a reporter assay to test whether overexpression of N regulates the expression of the main open reading frame following the 5′ UTR and stem loop 5 (SL5). To determine if the 5′ UTR or SL5 or both participate in any regulatory activity in concert with N, we inserted the SARS-CoV-2 5′ UTR with and without the complete SL5 upstream of GLuc (**Fig. 6a**). We also included a reporter with only the complete SL5 but not the rest of the 5′ UTR. The parent plasmid vector without any 5′ UTR inserted serves as a negative control and CLuc encoded on the same reporter plasmid serves as a transfection control. Thus, the ratio of GLuc to CLuc luciferase activity would measure the changes in translation as a result of overexpressing N (from a co-transfected plasmid), compared to a separate control co-transfection overexpressing mCherry. Our results show that overexpression of N leads to significantly lower expression of GLuc relative to CLuc, but only when the complete SL5 is present (pLT4 and pLT14 in **Fig. 6b**). The reporter containing only the full-length 5′ UTR, and therefore lacking the complete SL5, does not show differential expression. The change in GLuc/CLuc expression is not observed at the RNA level (**Fig. 6c**), which suggests that N regulates translation and not RNA stability. Given that N protein binding to this region significantly downregulates translation in our reporter assay, we sought to determine whether this effect results solely from binding to the 5′ UTR, possibly obstructing the translation initiation complex scanning. To test this, we used the Renilla-MS2/Firefly tethering assay to tether N to the 3′ UTR of Renilla-MS2 and found that it decreased Renilla luciferase activity (**Fig. 6d**), suggesting that other mechanisms of downregulation may be involved.

**Fig 6.**
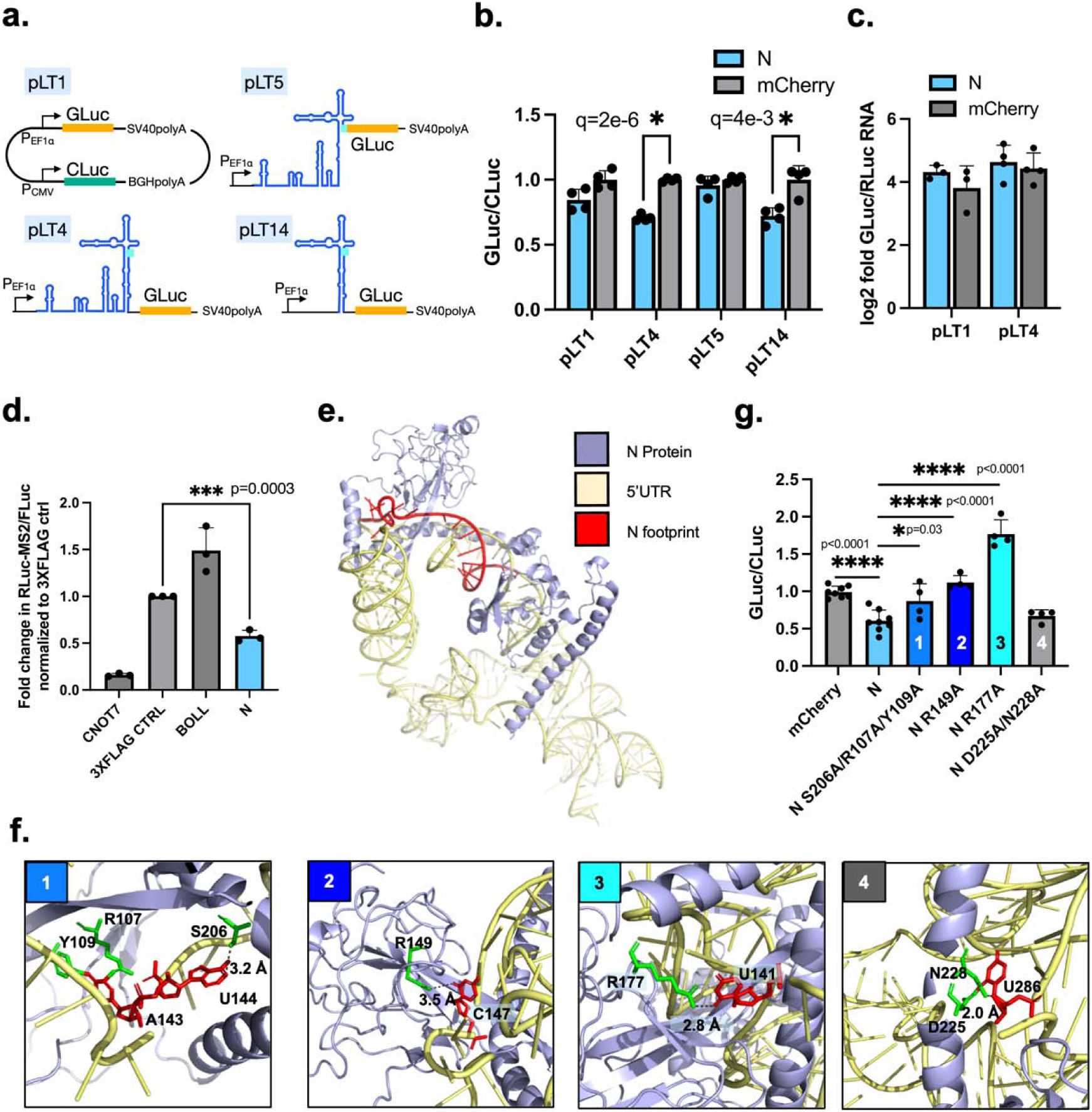
Interaction between N and 5′ untranslated region impacts translation. **a)** Schematic showing a dual luciferase reporter assay where different SARS-CoV-2 5′ end regions are inserted upstream of a Gaussia luciferase (GLuc) to regulate its expression. Cypridina luciferase (CLuc) serves as a transfection control in all four reporter constructs. pLT1 is a negative control vector with no 5′ UTR inserted. **b)** Luciferase activity GLuc/CLuc normalized to pLT1 when co-transfected with N, with mCherry as a negative control. *q values from multiple unpaired two-tailed t-test with Benjamini-Hochberg correction. **c)** Ratio of *CLuc* to *GLuc* RNA levels determined by RT-qPCR. **d)** Fold change in the RLuc/FLuc luciferase activity ratio for MCP-tagged proteins. 3×FLAG serves as a non-regulatory control, CNOT7 as a downregulatory control, and BOLL as an upregulatory control. *p-values from two-tailed unpaired t-tests. **e)** AlphaFold3 structure prediction of N and 5′ UTR-SL5 RNA. **f)** Magnified regions indicating N amino acid and nucleobase interactions. **g)** Relative GLuc/CLuc luciferase activity ratio due to overexpressing N mutants (corresponding to d) compared to wildtype N and mCherry. *p-values from two-tailed unpaired t-tests.

To further investigate the interactions between the N protein and the 5′ UTR, we modeled the structure of N bound to the 5′ UTR-SL5 RNA (nucleotides 1-294). Since there is no solved structure for the N protein and 5′ UTR-SL5 RNA complex, we used AlphaFold3 to predict the structure of the complex. As validation, the N-terminal domain (NTD) of the AlphaFold3-predicted structure aligned with the NTD of a solved crystal structure with a root-mean-square deviation (RMSD) of 0.373 Å, showing high structural similarity (**Supplementary Fig. 5**). From the predicted structure, we found that the N footprint RNA sequence forms numerous contacts (≤3.5 Å) with N, particularly within nucleotides 140-166 (**Fig. 6e-f, Supplementary Table 2**). These interactions involve positively charged residues (arginine and lysine) as well as polar side chains (serine, threonine, glutamine, and asparagine). To test whether these potential contacts contribute to N-mediated translational regulation, we generated individual mutations in amino acids predicted to contact RNA nucleobases. AlphaFold3 additionally predicted a few contacts outside the N footprint region, including an interaction between N D225/N228 and RNA U286, which we also tested. Mutations S206A, R107A, Y109A, R149A, and R177A resulted in less downregulation compared to wild-type N, whereas D225A/N228A showed no difference (**Fig. 6g**). We further confirmed that these mutations did not alter the stability of the proteins (**Supplementary Fig. 6**).

### Frameshift element regions regulate programmed ribosomal frameshifting in a N protein-dependent manner

Given our findings of N being associated with translation regulation at the 5′ UTR, we wondered if N is implicated in other translation events, such as programmed ribosomal frameshifting (PRF). SARS-CoV-2 encodes a ribosome frameshift region consisting of a slippery sequence (UUUAAAC) followed by a pseudoknot RNA structure, which causes the ribosome to move backward by one nucleotide before continuing translation (i.e. -1 PRF)^12^. The non-frameshifted polypeptide ORF1a ends with the stop codon after NSP11, whereas frameshifting begins the translation of NSP12 to continue synthesis of NSP13 through NSP16 to form the full-length polypeptide ORF1ab. We observed that N significantly enriched reads 5′ of the attenuator loop – a cis-regulatory element before the slippery sequence that dampens frameshifting – at region 13,415–13,435 nt with a mean fold enrichment of 3.3 (p<1e-24) (**Fig. 7a-b**). Neither NSP12 nor NSP8 show similar enrichment. Shannon entropy analysis indicates that this region is conserved among related betacoronaviruses (**Fig. 7c**). Intrigued by this conserved element, we hypothesized that N modulates ribosome frameshifting.

**Fig 7.**
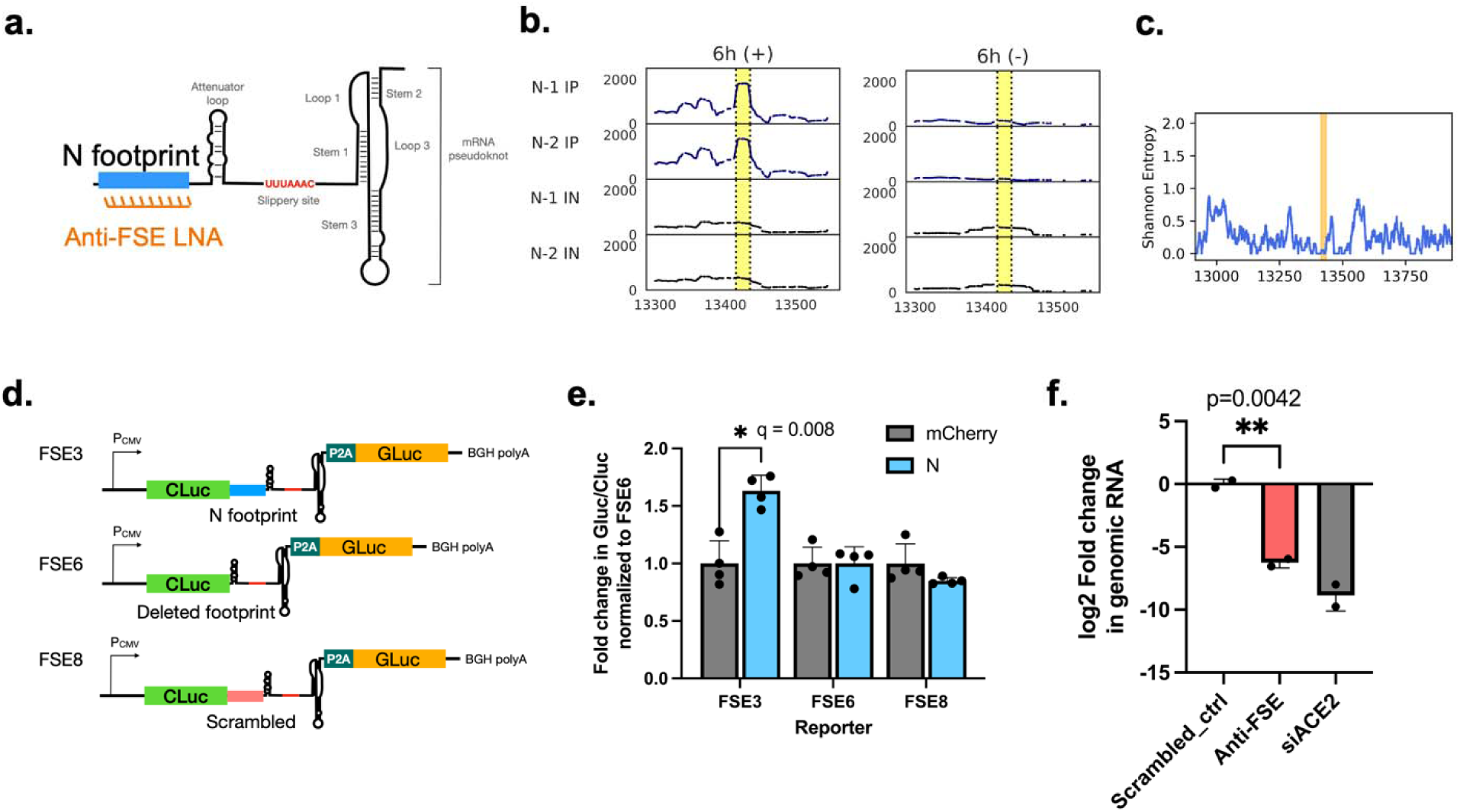
N binds upstream of the frameshift element and regulates ribosomal frameshifting. **a)** Schematic showing the frameshift element, attenuator loop, slippery site and pseudoknot, and highlighted in orange is a region that interacts with N, corresponding with the blue highlighted region in **b**. Anti-FSE LNA, locked nucleic acid targeting the N footprint. **b)** Read density plots showing RNA enriched by N eCLIP (N footprint). **c)** Shannon entropy zoomed into the region around the frameshift element; orange highlighted sequence indicates the 13,415 – 13,435 nt region bound by N. **d)** Schematic showing a dual luciferase reporter assay where the SARS-CoV-2 frameshift region is inserted upstream of a Gaussia luciferase (GLuc) to regulate its expression, with (FSE3), without (FSE6) the N footprint and with a scrambled region (FSE8). GLuc is -1 frameshifted from an upstream Cypridina luciferase (CLuc). GLuc is observed only if -1 frameshifting occurs. **e)** Luciferase activity GLuc/Cluc normalized to FSE6 when frameshift reporters are co-transfected with N. *q values from multiple Welch’s t-test with Benjamini-Hochberg correction. **f)** RT-qPCR results indicating log2 fold change of SARS-CoV-2 genomic RNA levels in virions released from cells treated with anti_FSE LNA, compared to a scrambled LNA control and siRNAs targeting ACE2. *p-values from two-tailed unpaired t-tests.

We developed a reporter assay incorporating the SARS-CoV-2 frameshift region to test whether overexpression of N leads to different degrees of frameshifting. We used a construct expressing GLuc and CLuc linked by the frameshift region, with GLuc out of frame from CLuc. GLuc is only expressed when -1 ribosome frameshifting occurs, thus the ratio of GLuc to CLuc luciferase activity measures the extent of frameshifting (**Fig. 7d**). We constructed three reporters – FSE3, FSE6 and FSE8: FSE3 contains the region bound by N (N footprint) in addition to the rest of the frameshift element; FSE6 does not contain the N footprint; and FSE8 contains a scrambled N footprint region (i.e. synonymous codons). Compared to FSE6 and mCherry as a negative control, N overexpression increases the ratio of GLuc to CLuc by approximately 50% (**Fig. 7e**). Our results show that overexpression of N leads to increased frameshifting when the N footprint is present.

To determine whether the N footprint at the FSE is essential to the virus, we designed a locked nucleic acid (anti-FSE_N_footprint LNA) that targets the region 13,415–13,435 nt (**Fig. 7a**). We hypothesized that disrupting any interaction of this region with other regulatory factors, including N, would exert an impact on virus proliferation. We treated cells with the anti-FSE_N_footprint LNA before infecting with live virus and observed a reduction in the amount of virus produced by the cells treated with this LNA compared to a Scrambled_LNA control (**Fig. 7f, Supplementary Fig. 3**). We conclude that the N footprint plays a significant and essential role in regulating frameshifting.

## Discussion

In this study, we used eCLIP to generate comprehensive, genome-wide maps of SARS-CoV-2 RNA interacting with NSP8, NSP12 and N at early and late stages of infection. Our analysis identified hundreds of regions with significant enrichment by each of the proteins. NSP12 and NSP8 eCLIP results strongly enrich RNA reads at the 5′ and 3′ untranslated regions (UTRs) of both the positive and negative sense virus RNA. NSP12 and NSP8 interact extensively with negative-strand RNA during early infection but show little to no interaction at late stages. This pattern reflects their role in supporting the high levels of replication and transcription that occur early in infection. The absence of interaction with the negative strand at later time points may not necessarily indicate reduced affinity for negative-strand RNA, but rather the accumulation of more positive strand RNA due to a transcriptional bias favoring the production of positive-strand RNA^28^ (**Fig. 4f**). Additionally, we observed widespread interaction between N and only the positive sense viral RNA at both early and late stages of infection. The number of interactions between N and viral RNA increases substantially from early to late infection, which supports our understanding of its role in condensing genomic RNA to facilitate assembly and packaging of virions^19^. While this manuscript focuses on the gene regulatory potential of viral protein–RNA interactions, our dataset offers a rich resource for future investigations into the mechanisms of viral genome assembly.

We identified a novel regulatory element within the RNA sequence that encodes the highly conserved but poorly understood Y1 domain of NSP3. Our findings show that this element, bound by NSP12, regulates and coordinates the RNA stability or translation of sequences upstream of the NSP12 footprint. As evidence of different levels of viral RNA upstream of the NSP12 binding site, the drop off in RNA reads extending 0.5-1.5 kilobases from the 5’ end of the genome has also been shown by others^27^. Notably, targeting the NSP12 footprint sequence with an antisense locked nucleic acid significantly impacted viral fitness, suggesting this conserved region to play a critical role in the virus life cycle. Thus, NSP12 contributes to regulating the RNA levels of viral proteins, though the precise mechanism requires further investigation. Coronaviruses commonly undergo genomic recombination to increase genetic diversity and host adaptability. Recombination occurs when the virus RNA-dependent RNA polymerase (RdRp) is dissociated from one RNA template before reassociating and continuing replication on a different template^31,32^. Recombination of SARS-CoV-2 strains near the NSP12 footprint has been observed in an XAY strain, which is a result of recombination between the Delta strain and the Omicron BA.2 strain^33^. This may be linked to NSP12 interactions near the recombination breakpoint. The structured region at the NSP12 footprint may stall the polymerase during elongation, causing it to dissociate and resulting in reduced levels of downstream RNA. Additionally, NSP12 has been reported to affect translation elongation of host RNA^14^. However, much more work is needed to elucidate the underlying mechanism. Our results identify an essential and complex role of the Y1 coding RNA and its interaction with NSP12 in the virus life cycle.

Furthermore, we found a novel regulatory site at the 5’ UTR bound by N since early stages of infection in virus infected cells. A recent study used NMR spectroscopy to characterize the low micromolar affinity binding of the N terminal domain of N protein and stem loop 5^34^. Our study provides evidence of this interaction in authentic virus infected cells. The study showed R107A to be a binding deficient mutant, which strengthens our findings that the interaction of this amino acid with the 5’ UTR affects the regulatory activity of N. We have further shown that this binding site is essential to the virus via LNA targeting. Since the 5’ UTR plays a crucial role in translation initiation regulation, we hypothesized that N binding to this region could be regulating virus translation. Additionally, we identified an N protein interaction at the frameshift region, demonstrating complex involvement of N in regulating translational processes. The propensity for N to be implicated in regulating translation is supported by literature evidence. It has been found to potentiate host NPM1-snoRNA translation machinery to enhance viral replication^35^. N is also bound to PABPC1/4^1^, which binds to eIF4G and eIF4E to facilitate translation initiation and it was shown in BCoV that it may be involved in translation inhibition. Downregulating viral translation may be a strategy employed by the virus to switch from an early infection stage with high translation initiation to a later stage of low translation as genome assembly and viral packaging become primary processes. Since the N protein binds to SL5, which is located downstream of the leader peptide that precedes subgenomic RNAs, its downregulation of ORF1ab translation could help redirect resources toward the translation of subgenomic RNA-encoded structural and accessory proteins.

A limitation of our study is that the eCLIP data is an aggregate of all the binding events taking place in the whole cell. However, the viral RNA is compartmentalized for RNA synthesis in the double-membraned vesicles (DMVs) of replication^36^ before subsequently released into the ER-Golgi intermediate compartment (ERGIC) for translation and genome assembly with N. This spatial separation imposes constraints on interpretation. For example, our data do not distinguish whether interactions between NSP12/NSP8 and the 5′ UTR occur within replication centers, where they may regulate RNA synthesis, or in the ERGIC, where such interactions could affect translation or genome packaging. Similarly, NSP12 binding to the Y1 coding region may not influence translation if the interaction occurs inside DMVs, unless NSP12 remains bound to the RNA as it is transported into the cytoplasm. Notably, because this binding is observed on both positive- and negative-sense strands, it may reflect interactions with double-stranded RNA intermediates in DMVs. Without single-molecule resolution or isoform specificity, our eCLIP data cannot distinguish between subgenomic mRNAs and the full-length positive-sense genome. The observed preference of N for positive-sense RNA may reflect either the sequestration of negative-strand RNA within replication vesicles or selective binding to sequence or structural features. Approaches that can capture these spatial and isoform differences will be needed to address these limitations.

In conclusion, our genome-wide interrogation of the interaction of SARS-CoV-2 NSP8, NSP12 and N proteins with viral RNA not only sheds light on known regulatory regions, but also uncovers new sites previously unknown to play a role in virus fitness. Since targeting these regions using LNAs can reduce viral fitness, these interaction sites may serve as potential drug targets, which could open up possibilities for future antiviral drug development. RNA structures and RNA-protein interaction sites are emerging as a new class of druggable targets, which will substantially expand our current capabilities in antiviral drug discovery. Given the conservation of the regulatory regions we identified, insights from SARS-CoV-2 may help inform strategies to safeguard against future outbreaks of related SARS-like viruses. Moreover, since many RNA viruses encode functionally N-like proteins, our findings may have broader relevance across other viral families.

## Methods

### Cell culture

Vero E6 cells and HEK293T cells were purchased from the American Type Culture Collection and were not further authenticated. Cells were routinely tested for mycoplasma contamination with a MycoAlert mycoplasma test kit (Lonza) and were found negative for mycoplasma. Growth media was replaced every two days, and the cells were passaged every four days. HEK293T and Vero E6 cells were cultured in DMEM (ThermoFisher) supplemented with 10% FBS (ThermoFisher) and passaged every three days. All cell cultures were incubated at 37°C and 5% CO_2_.

### SARS-CoV-2 virus infection

Infectious SARS-CoV-2 was conducted in Biosafety Level-3 conditions at the University of California San Diego and the University of California Riverside following the guidelines approved by the respective Institutional Biosafety Committees. SARS-CoV-2 isolates USA-WA1/2020 (BEI Resources, #NR-52281), hCoV-19/USA/CA_UCSD_5574/2020 (lineage B.1.1.7) and hCoV-19/South Africa/KRISP-K005325/2020 (lineage B.1.351 BEI Resources NR-54009) were propagated and infectious units quantified by plaque assays and fluorescent focus assay using TMPRSS2-Vero E6 cells (Sekisui XenoTech). Viral stocks were confirmed by whole genome sequencing.

For eCLIP assays, Vero E6 cells were seeded at 5 million cells 24 hours before infection. About an hour before infection, the culture media was changed from 10% FBS to 2% FBS in DMEM. Cells were infected at a multiplicity of infection (MOI) of 1 and incubated for 6 hours. Additionally, cells were infected at a MOI of 0.01 and incubated for 48 hours. Infected cells were then rinsed with 1XPBS and a thin layer of chilled PBS was then added. Cells were crosslinked on a chilled metal block in a UVP Crosslinker CL-3000 (Analytik Jena) with UV_254nm_ 400 mJ/cm^2^. After crosslinking, the plates were removed, and the cells were scraped manually and spun down at 300 x g for 3 min at 4 degrees C. The supernatant was discarded, and the pelleted cells were kept at 4°C for transfer to the BSL2 laboratory to be snap frozen and stored at -80°C until ready for eCLIP processing.

### eCLIP library preparation and sequencing

The eCLIP experiment was performed as previously described^17^. Cell pellets were lysed using iCLIP buffer and lysates were sonicated and treated with RNase I to fragment RNA. Each sample of cell lysate was immunoprecipitated using 10 µg of primary antibodies that bind Nucleocapsid, NSP12 and NSP8: SARS-CoV / SARS-CoV-2 (COVID-19) NSP8 antibody Mouse monoclonal [5A10] (GeneTex GTX632696), SARS-CoV-2 (COVID-19) RdRp (nsp12) antibody rabbit polyclonal (GeneTex GTX135467), SARS-CoV-2 (2019-nCoV) Nucleocapsid rabbit polyclonal (Sino Biological 40143-R019). Primary antibodies bound to Sheep Anti-Mouse IgG Dynabeads M-280 (ThermoFisher) were incubated with lysates overnight at 4°C. Two percent volume of each lysate sample was stored for preparation of a parallel SMInput library. The remaining lysates were processed as the immunoprecipitated (IP) samples. High salt wash buffer containing 50 mM Tris-HCl pH 7.4, 1% Tween-20, 1 M NaCl, 1 mM EDTA 1M HCl, 1% NP-40, 0.1% SDS 1M NaOH, 0.5% sodium deoxycholate was used to remove most protein-protein interactions, as per standard eCLIP protocol. Bound RNA fragments in the immunoprecipitates were dephosphorylated and 3 -end ligated to an RNA adaptor. Protein– RNA complexes from SMInputs and immunoprecipitates were run on 4-12% BisTris SDS– polyacrylamide gel and transferred to nitrocellulose membrane. Membrane regions comprising the exact protein sizes to 75 kDa above were excised, and RNA was released from the complexes with proteinase K. SMInput samples were dephosphorylated and 3 -end ligated to an RNA adaptor. All RNA samples (immunoprecipitates and SMInputs) were reverse transcribed with SuperScript III Reverse Transcriptase (LifeTech 18080044). cDNAs were 5 -end ligated to a DNA adaptor. cDNA yields were quantified by qPCR, and 100–500 fmol of library was generated with Q5 PCR mix (NEB). Library sequencing was performed at the UCSD IGM Genomics Center.

### Analysis of eCLIP sequencing data

For eCLIP performed on SARS-CoV-2 infected Vero E6 cells, analysis was adapted from a published eCLIP pipeline (https://github.com/yeolab). Sequencing reads were adapter-trimmed and mapped to SARS-CoV-2 MN908947.3 genome assembly. PCR duplicate reads were removed using the unique molecular identifier sequences in the 5 adaptor, and remaining reads were retained as ‘usable reads’. For reads mapped to the SARS-CoV-2 genome, bedgraph densities were generated using SAM tools v1.9 to obtain read densities at each nucleotide position. To determine regions with significant enrichment by immunoprecipitation, read densities mapped to the virus genome are analyzed in windows of 20 nucleotides (**Supplementary Data**). Differences in read densities in each window of IP and SMInput samples are compared to determine the fold enrichment. Significance was determined using two-tailed unpaired t-test and false discovery rate q values was computed from t-test p-values using the Benjamini–Hochberg procedure. Windows with log2foldchange >1.5, q-value < 0.01 were designated as significant. Specific nucleotide regions for manual examination were determined using two-tailed unpaired t-test.

### LNA design, transfection, and SARS-CoV-2 infection

Antisense locked nucleic acids (LNAs, Integrated DNA Technologies) were designed to anneal to target sequences within the SARS-CoV-2 genome (GenBank: MN908947), similar to previously reported^11^. Briefly, all LNAs were designed with three consecutive LNA bases at the 5′ and 3′ ends of each oligonucleotide, with stretches of unlocked bases within the oligonucleotide limited to three consecutive nucleotides. All LNAs were designed with similar thermodynamic properties, including length, %GC content, %LNA content, and LNA:RNA duplex T_m_ (**Supplementary Table 1**).

Vero E6 cells were plated at 200,000 cells per well of a 12-well plate. LNAs were transfected at a final concentration of 400nM per well using Lipofectamine 2000. siRNAs targeting ACE2 (hs.Ri.ACE2.13.1 and hs.Ri.ACE2.13.2 from Integrated DNA Technologies, 1:1 mixture of 5nM each) were used as a positive knockdown control. Cell viability of LNA transfected Vero E6 cells was determined by the CellTiter-Glo 2.0 Cell Viability Assay (Promega G9241) and measured on a Molecular Devices SpectraMax® iD5 Multi-Mode Microplate Reader and cytotoxicity was not observed (**Supplementary Figure 4**). One day after transfection of LNAs and siRNAs, the growth media was replaced with serum free DMEM. Transfected cells were infected by SARS-CoV-2 at an MOI of 0.01 for 24 hours. For downstream RNA extraction and RT-qPCR, virus was inactivated with RLT buffer for 15 min before transferring from BSL3 to BSL2. RNA extraction was performed using the Zymo Direct-zol RNA miniprep kit (R2052). RT-qPCR to quantify viral RNA encoding the Nucleocapsid protein was performed using iScript cDNA Synthesis Kit (Bio-Rad 1708891), iTaq SYBR® Universal Green Supermix qPCR Mastermix (Bio-Rad 1725125). Primers for RT-qPCR are SARS-CoV2_IBS_N1 forward CAATGCTGCAATCGTGCTAC and reverse GTTGCGACTACGTGATGAGG^38^, and readings were performed on a Bio-Rad CFX real-time system. Data analysis of Cq values extracted from Bio-Rad CFX software was performed using Graphpad Prism 10.

### Multiple sequence alignment and phylogenetic analysis

Complete genomes of betacoronavirus sequences from NCBI and GISAID were downloaded in March 2025. Multiple sequence alignment was performed using MAFFT v7.453 and default parameters of the local alignment l-ins-i. The phylogenetic tree was constructed using the average distance algorithm from the multiple sequence alignment and visualized within Jalview (version 2.11.4.1). Shannon entropy was calculated based on the log odds ratio of the base composition. H(n) = - Σ p(x) * log2(p(x)), where H(n) is the entropy at position n, p(x): is the probability of a particular nucleotide {A, U, G, C}, Σ: represents the summation over all four nucleotides.

### Dual luciferase plasmid construction and activity assays

Plasmids for reporter assays were derived from a dual luciferase reporter system based on ref^39^ (Addgene #181934). It encodes *Cypridinia* and *Gaussia* luciferases (CLuc and GLuc) under the control of bidirectional EF1a and CMV promoters, respectively. Briefly, pLT1 was generated from this plasmid by addition of StuI and EcoRI restriction sites at in the 5’ UTR and XbaI and BamHI restriction sites before and after the SV40 polyA sequence in the 3’ UTR. SARS-CoV-2 5’ UTR and stem loop 5 reporters (pLT4, pLT5 and pLT14) were generated by insertion of 5’ UTR and stem loop 5 using oligonucleotides (IDT) or gene fragments (GenScript) at the 5’ end of GLuc. Frameshift reporters FSE3, FSE6 and FSE8 and Y1 reporters were assembled using fragments that are PCR amplified from the pLT1 plasmid such that CLuc and GLuc are transcribed using the same EF1α promoter, and gene fragments were synthesized by GenScript. Plasmids encoding NSP12 and N were subcloned from ref^1^ (Addgene #141391 and #141378) into a pcDNA3.4 vector. The authors will deposit these plasmids on the Addgene repository to be made publicly available before publication.

Luciferase assays were performed via co-transfection of dual reporter plasmids with plasmids expressing NSP12, N and a pcDNA3.1 plasmid expressing mCherry^40^ using Lipofectamine 2000 reagent (Invitrogen 11668019) and according to manufacturer’s protocol. Luciferase activity assays were performed using the UltraBrite Cypridina-Gaussia dual luciferase assay reagent from Targeting Systems (DLAR-4 SG-1000) and measured on a Molecular Devices SpectraMax® iD5 Multi-Mode Microplate Reader using an integration time of 2 seconds/well. Data analysis was performed using Graphpad Prism 10.

### Western blot

Cells were washed with PBS and lysed in RIPA buffer with Protease Inhibitor Cocktail Set III (EMD Millipore). Lysates were homogenized using Qiashredder (Qiagen). Protein extracts were denatured at 75 °C for 20 min and run at 150 V for 1.5 h on 4-12% NuPAGE Bis-Tris gels in NuPAGE MOPS running buffer (Thermo Fisher). Proteins were transferred to polyvinylidene difluoride membrane using NuPAGE transfer buffer (Thermo Fisher) with 10% methanol. Membranes were blocked in blocking buffer (TBS containing 5% (wt/vol) dry milk powder) for 30 min and probed with primary antibodies in blocking buffer for 16 h at 4 °C. Membranes were washed three times with TBST and probed with secondary HRP-conjugated antibodies in blocking buffer for 1 h at room temperature. Signal was detected by Biorad ECL and imaged on an Azure Biosystems C600 imager, or UVP ChemSolo imager (Analytik Jena).

### Data availability

Plasmids and cell lines generated in this work are available upon request. All sequencing data are deposited in GEO with accession GSE173508.

## Acknowledgements

We appreciate Dr. Roya Zandi and Dr. Siyu Li, and members of the Xiang and Yeo lab for providing helpful discussions. We thank the Nikon Imaging Center, University of California, San Diego, for imaging assistance. The work has been supported by Emergency COVID-19 Research Seed Funding (#R00RG2636) from the University of California Office of the President. This publication includes data generated at the UC San Diego IGM Genomics Center utilizing an Illumina NovaSeq 6000 that was purchased with funding from a National Institutes of Health SIG grant (#S10 OD026929). AFC is supported by an NIH grant K08 AI130381 and a Burroughs Wellcome Fund Career Award for Medical Scientists. The following reagents were deposited by the Centers for Disease Control and Prevention and obtained through BEI Resources, NIAID, NIH: SARS-Related Coronavirus 2, Isolate USA-WA1/2020, NR-52281. The following reagent was obtained through BEI Resources, NIAID, NIH: SARS-Related Coronavirus 2, Isolate hCoV-19/South Africa/KRISP-K005325/2020, NR-54009, contributed by Alex Sigal and Tulio de Oliveira.

## Author contributions

J.S.X. and G.W.Y. conceived of the project. J.S.X, K.X.Z, J.R.M, J.C.S, K.R., R.N.M, B.A.C., A.F.C. and S.L.L designed and performed experiments. L.T., K.T., S.S.P, E.M.K, A.A.M., B.A.C, A.E.C., and C.A. performed experiments. J.S.X, J.R.M, J.C.S, K.R., A.F.C., S.O’L., R.H., S.L.L analyzed experimental results. J.S.X and B.A.Y performed bioinformatics analysis. J.S.X, S.L.L and G.W.Y wrote the manuscript with help from all authors. J.S.X and G.W.Y supervised the project.

