## Supplementary Information for "Genome-Wide Interrogation of SARS-CoV-2 RNA-Protein Interactions Uncovers Hidden Regulatory Sites"

### Supplementary Fig. 1.

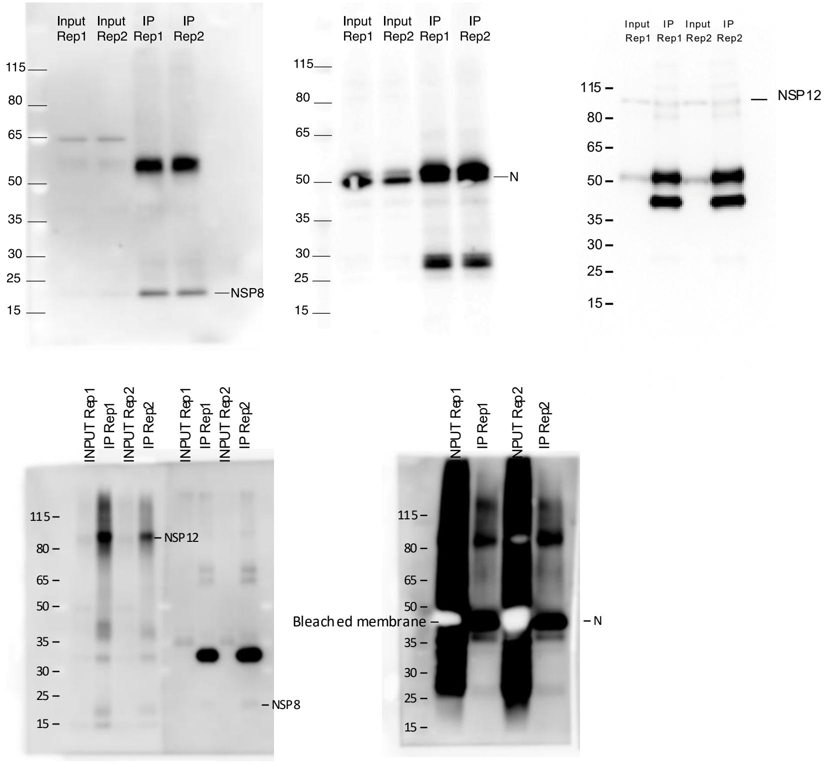


**Supplementary Fig. 1.** Western blots indicating the presence of input and immunoprecipitated proteins NSP8, NSP12 and nucleocapsid at 6 hours (a) and 48 hours post infection (b).

#### Supplementary Fig. 2.

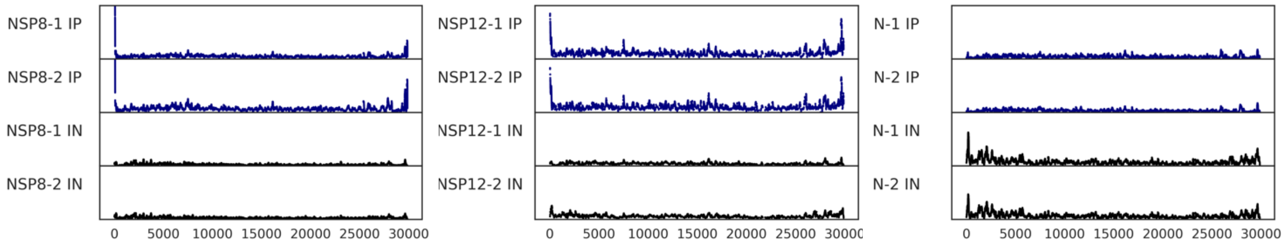


**Supplementary Fig. 2.** Read density plots show RNA enriched by NSP8, NSP12, and N distributed across the negative same SARS-CoV-2 genome at 6 hours post infection. Two biological replicates were performed for each sample and condition.

**Supplementary Fig. 3.**

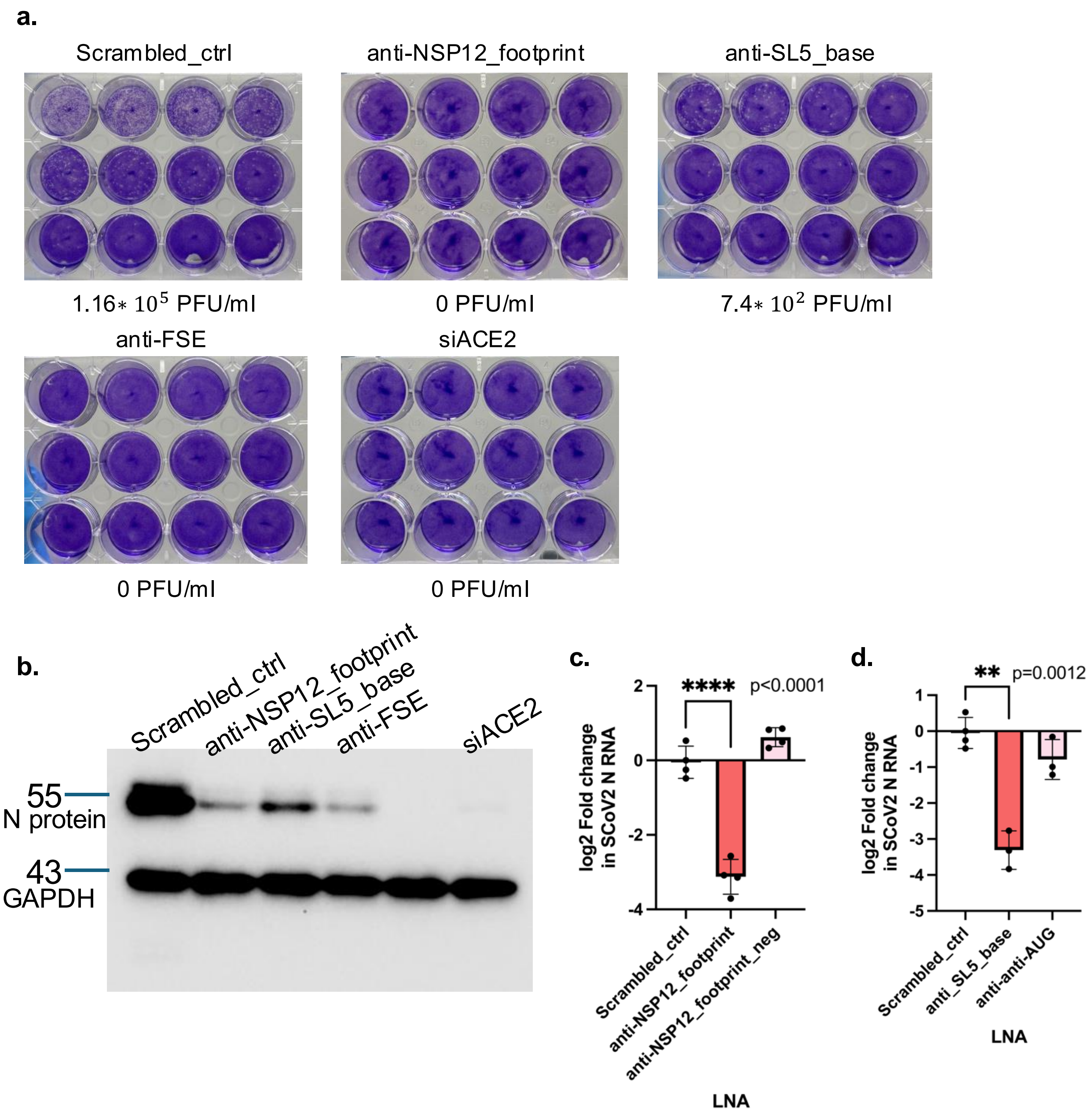


**Supplementary Fig. 3. a.** Viral titer of Vero E6 cells treated with LNA prior to infection was quantified using plaque assay. Each well was infected with designated dilutions of SARS-CoV-2 infected and LNA treated samples ranging from $10^{-2}$ to $10^{-5}$. **b.** Western blot analysis on the amount of N protein in Vero E6 cells treated with LNA prior to infection. Cells were infected at an MOI of 0.01 for 24 hours in all samples. RT-qPCR results indicating log2 fold change of SARS-CoV-2 N RNA levels in cells treated with LNAs that target the Y1 NSP12 footprint region (**c**) and the SL5 base N footprint region (**d**). P-values are two-tailed Welch's t-test.

**Supplementary Fig. 4.**

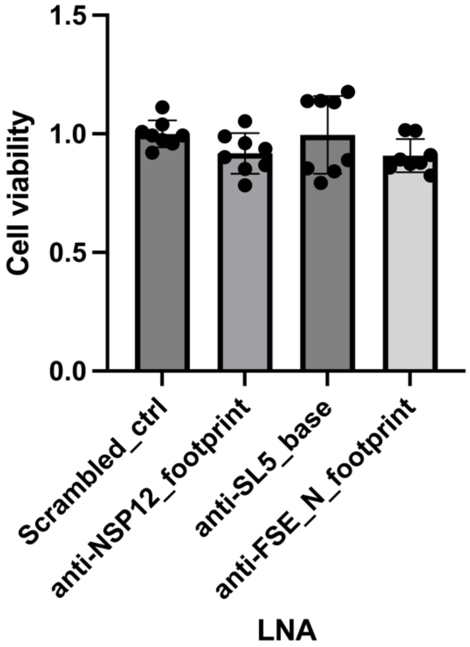


**Supplementary Fig. 4.** Cell viability assay of Vero E6 cells transfected with LNA. Cell viability is given by luciferase activity normalized to cells transfected with the Scrambled_ctrl LNA. N=8 biological replicates.

**Supplementary Fig. 5.**

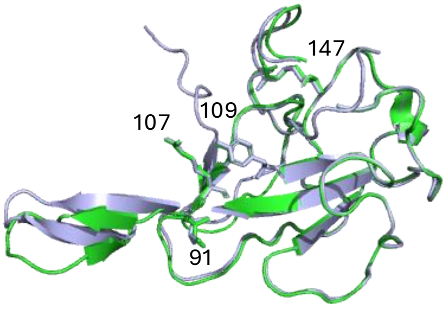


**Supplementary Fig. 5.** Superposition of the N-terminal domain crystal structure (PDB ID: 9EXB, green) with the NTD of the AlphaFold3 predicted structure used in Main Figure 6 (purple). RMSD of the alignment is 0.373 Å.

**Supplementary Fig. 6.**

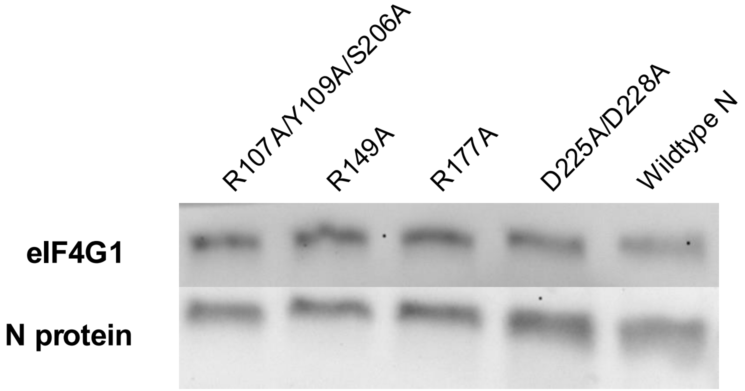


**Supplementary Fig. 6.** Western blot of N mutants and wildtype N. eIF4G1 is used as a housekeeping gene.

**Supplementary Table 1.** Locked nucleic acids (LNAs).
+{A,C,T,G} refers to the locked nucleic acid version of the nucleotide
Melting temperature Tm is predicted using IDT's OligoAnalyzer
Region_15, Region_15_ctrl, Region_22 and Region_22_ctrl are shown below as reference.

| LNA Name | Sequence | Tm (ºC) | Source |
|---|---|---|---|
| Scrambled_ctrl | +G+C+GGC+ACG+TTG+CG+AGT+A+C+T | 72.8 | Huston et al |
| Region_15 | +A+C+AAA+CCC+TTG+CCG+AG+CT+G+C+T | 77.3 ºC | Huston et al |
| Region_15_ctrl | +G+T+TT+TCA+ACT+TTG+TTA+TAG+G+T+G | 62.3 ºC | Huston et al |
| Region_22 | +G+T+CTA+ACA+ACA+TCA+AA+AG+G+T+G | 66 ºC | Huston et al |
| Region_22_ctrl | +G+C+TAC+AG+TGG+CAA+GAG+AA+G+G+T | 71.7 ºC | Huston et al |
| Anti_NSP12_footprint | +T+A+ACA+CAT+CAT+ACA+AGT+TG+AT+G+A+A | 62.7 | This work |
| Anti_NSP12_footprint_neg | +T+A+CAA+GTT+GAT+GAA+TTA+CA+A+C+C | 57.7 | This work |
| Anti_SL5_base | +T+C+CTG+TCA+ACG+AC+AG+TAA+T+T+A | 61.2 | This work |
| Anti_anti_AUG | +G+A+TAG+ACG+AGT+TAC+TCG+TGT+C+C+T | 68.3 | This work |
| Anti_FSE | +G+C+GG+AGT+TGA+TCA+CA+A+C+T | 65.3 | This work |

**Supplementary Table 2.**Table listing AlphaFold3 predicted interaction sites within the N footprint.

| 5′ RNA position | 5′ RNA base | N position | N amino acid | Distance (Å) | RNA Contact |
|---|---|---|---|---|---|
| 153/154 | U/U | 261 | K | 2.7/3.2 | backbone |
| 164/165 | C/G | 303 | Q | 2.5/2.2 | backbone |
| 163 | A | 299 | K | 2.5 | backbone |
| 160/161 | G/A | 240 | Q | 0 | backbone |
| 166 | A | 345 | N | 0 | backbone |
| 149/150 | G/U | 49/88 | T/R | 2.6/2.8/3.5 | backbone |
| 147 | C | 149 | R | 3.5 | base |
| 143/144 | A/U | 206/ (107/109) | S/(R/Y) | 3.2/(0/0) | base |
| 140 | C | 209 | R | 3.5 | backbone |
| 141 | U | 177 | R | 2.8 | base |
| 148 | U | 49/111 | T/Y | 2.7/3.1 | backbone |
